# De Novo Design and AlphaFold3 Evaluation of Protein Binders Targeting Specific Sites of MAP4K4

**DOI:** 10.64898/2026.09.04.749196

**Authors:** Aarnav Jain, Alexander V. Tobias

## Abstract

MAP4K4 is a serine/threonine kinase member of the mitogen activated protein kinase family. It acts through the JNK, p38 MAPK, and ERK1/2 pathways and is implicated in cancer proliferation and invasion, TNF-α-driven insulin resistance, macrophage-mediated inflammation, and cardiomyocyte apoptosis in heart failure. No selective small-molecule inhibitor of MAP4K4 has reached clinical use, and knowledge of the specific epitopes on MAP4K4 is lacking for some existing antibodies, leaving a role for small protein binders directed at specific surface sites. We used a computational pipeline that combines RFdiffusion, ProteinMPNN, and AlphaFold2, as well as BindCraft, an integrated “one-shot” tool, for *de novo* design of small protein binders (∼50–130 residues) to specific MAP4K4 surface hotspots. We also created an automated hotspot determination algorithm that weighs geometry, chemistry, rigidity, and AlphaFold pLDDT. From thousands of candidate sequences, we evaluated 20 of the most promising binders with AlphaFold3 (AF3). The interface predicted template modeling (ipTM) scores ranged from 0.16 to 0.90, with nine candidates having ipTM ≥ 0.80, and five scoring ≥ 0.87. Two binders engage non-overlapping hotspots on opposite faces of MAP4K4, making them a candidate pair for a sandwich assay. BLASTp searches of all designed protein sequences returned only low-significance matches to half of them, indicating that they represent truly novel binding solutions and previously unexplored regions of protein sequence space rather than rediscovered natural motifs. We subjected five complexes spanning the observed AF3 confidence range to 100-ns explicit-solvent molecular dynamics simulation. Interchain contacts were retained throughout, with stability varying substantially between systems. Confidence in the binding specificity of the nine highest-confidence candidates was bolstered by juxtaposition with AF3 evaluations of their interaction with CDK2, a negative-control kinase. This comparison yielded a significant, consistent reduction in ipTM (p = 0.0039) for the control binding partner. ToxinPred2 and AlgPred 2.0 screening suggested that one candidate was a potential allergen and two were potential toxins. These results support that *de novo* design of small, site-specific protein probes for an underserved disease target is achievable using free, publicly available computational tools and minimal resources, pointing to a greater role for the public and amateur scientists to contribute to biotechnological advancement.

## 1. Introduction

MAP4K4 (mitogen-activated protein kinase kinase kinase kinase 4) is a serine/threonine kinase that signals through the JNK, p38 MAPK, and ERK1/2 cascades. Research has suggested that MAP4K4 is a part of several major disease processes. It is overexpressed across colorectal, gastric, pancreatic, lung, and hepatocellular cancers, where higher expression correlates with poor prognosis, invasion, and metastasis [1]. In adipocytes and skeletal muscle, TNF-α signaling upregulates MAP4K4, which in turn suppresses insulin-stimulated glucose uptake and drives insulin resistance [2]. In macrophages, MAP4K4 mediates TNF-α and IL-1β production, and silencing it *in vivo* protects mice from LPS-induced inflammatory lethality [3]. MAP4K4-deficient T cells stabilize TRAF2 and preferentially differentiate into pathogenic Th17 cells, linking loss of MAP4K4 activity to chronic inflammation [4]. In the heart, MAP4K4 activity rises in failing human myocardium and drives cardiomyocyte apoptosis through the TAK1-JNK pathway [5]. Despite this broad involvement in cancer, metabolic disease, inflammation, and heart failure, no selective small-molecule inhibitor of MAP4K4 has reached clinical use. Generally speaking, knowledge of the specific epitope for many antibodies against protein targets is lacking [6]. This leaves the surface of MAP4K4 largely unexplored as a target for engineered binders directed toward specific sites.

Recent advances in artificial intelligence have made *de novo* protein design practical without large-scale wet-lab design and screening. Diffusion-based models can generate novel protein backbones with a specified shape or binding geometry (RFdiffusion [7]); deep learning sequence-design models can then assign an amino acid sequence expected to fold into that backbone (ProteinMPNN [8]); and structure-prediction models can evaluate whether a designed sequence actually adopts the intended fold and engages the intended target (AlphaFold2 [9]). BindCraft [10] packages this functionality into a more self-directed and checkpoint-governed one-shot design pipeline, further reducing the computational and expertise barrier to generating candidate binders. Because both pipelines’ internal AlphaFold 2 (AF2)-based confidence scores can be overly optimistic, subsequent evaluation is an important check. The AlphaFold Server (AFS; alphafoldserver.com) running AlphaFold3 (AF3) [11] provides a publicly accessible way to assess a candidate complex outside of these design pipelines. AF3 offers several advances over AF2. These include an architecture that models binder-target complexes with a generalized atomic-level representation, a generative diffusion model that directly predicts physical atom coordinates without the need for complex physics-based constraints, and reduced reliance on multiple sequence alignments with improved accuracy for multi-chain assemblies like noncovalent protein complexes [12]. DeepMind initially withheld AF3’s code and model weights, due to biosecurity concerns [13], however both have since been released as an inference pipeline via GitHub, with weights available from DeepMind on request. Notwithstanding, integrating AF3 directly into BindCraft or RFdiffusion/ProteinMPNN was highly impractical for this project. We therefore used the pipelines with minimal modification to generate candidate binders (see Materials and Methods), followed by evaluation of these hits using the AFS web interface.

In this study, we combine an automated MAP4K4 surface-hotspot scoring heuristic with two parallel *de novo* design routes: a pipeline combining RFdiffusion, ProteinMPNN and AlphaFold2 (“RMA2” pipeline), and the BindCraft integrated suite. We subjected every top-ranked variant to independent scoring by AF3, and consider only sequences that scored well by AF3 to be high-confidence binder candidates. We report the sequences and evaluation scores a set of AF3-tested candidate MAP4K4 binders, compare AF2-reported and AF3-generated confidence scores to evaluate the extent of their agreement, characterized the predicted interface contacts of the strongest candidates, and assessed the sequence novelty of all designs with Basic Local Alignment Search Tool (BLAST) [14,15] queries. We also probed binding partner specificity with an AF3-based negative-control screen against an unrelated kinase. For five candidates ranging in binding interface scores as assessed by AF3, we additionally evaluated predicted binding-pose stability with explicit-solvent molecular dynamics (MD) simulation.

## 2. Materials and Methods

### 2.1 Experimental and Technical Design

Figure 1 summarizes the overall experimental design. Candidate MAP4K4 binders were generated via one of two *de novo* design pipelines implemented in the Google Colaboratory cloud-based programming environment: RFdiffusion/ProteinMPNN/AlphaFold2 [16] or BindCraft [10], both seeded from the same automatically detected surface hotspots on the MAP4K4 structure (PDB ID: 4U40). Every top-ranked candidate from either pipeline was independently evaluated with AlphaFold3 (AF3) on the AlphaFold Server before being reported herein. High-confidence binders were visualized with ChimeraX software [17] and screened for sequence novelty by BLAST.

**Figure 1.**
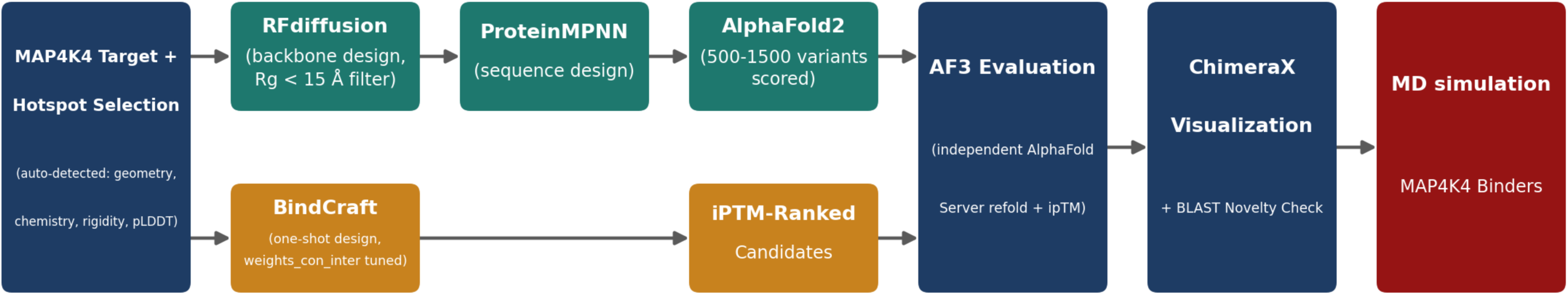
Computational pipeline for *de novo* design and AF3 evaluation of MAP4K4 binders. Candidate binders were generated via two parallel routes: RFdiffusion-generated backbones filtered by radius of gyration (R_g_ < 15 Å), followed by ProteinMPNN sequence design and AlphaFold2 scoring of 500-1500 resampled variants (top); and BindCraft one-shot design with specially tuned parameters (Section 2.4) to anchor designs at the intended hotspot (bottom). Both pipelines were fed hotspots from the same auto-detected MAP4K4 surface hotspots. Top-ranked candidates from either route were independently evaluated with AlphaFold3 before acceptance, then visualized with ChimeraX and searched by BLAST for novelty.

### 2.2 Target structure and hotspot selection

The MAP4K4 structure used as the design target for both pipelines was chain A of PDB entry 4U40 [18]. Candidate surface hotspots were identified with a custom coded automated scoring tool. Each surface residue was scored on four weighted criteria: local surface geometry (35%; e.g., grooves and concave patches versus flat surface), local chemistry (35%; hydrophobic character with a surrounding polar rim), backbone rigidity (20%; favoring residues in stable secondary structure over flexible loops), and AlphaFold’s predicted local distance difference test (pLDDT; 10%). Top-scoring residues were then clustered into contiguous 3-5-residue patches specified as input “hotspots” for either design pipeline.

### 2.3 De novo design via the RFdiffusion-ProteinMPNN-AF2 (RMA2) pipeline

For each selected hotspot, RFdiffusion [7] was used to generate novel binder backbones targeted to that site. Generated backbones were filtered by radius of gyration (R_g_ < 15 Å) to favor compact, globular designs before proceeding to sequence design. ProteinMPNN [8] was then used to assign amino acid sequences to each retained backbone. For simplicity, cysteine was excluded from the amino acid repertoire of ProteinMPNN. Designed sequences were evaluated with AF2 in-pipeline [9]. Certain backbones with high confidence results were subjected to additional sequence diversification with ProteinMPNN, producing hundreds of additional “second-generation” sequences evaluated by AF2 in search of superior scores, particularly interface predicted template modeling (ipTM). This truncated pipeline is referred to as “PA2.” This second-generation diversification step was applied only to the RMA2 pipeline (not to BindCraft) and, across all hotspots, produced sequences that were scored across 500–1500 resampled variants per retained backbone.

### 2.4 De novo design via the BindCraft pipeline

BindCraft [10], a distinct design pipeline was run against the same set of hotspots. Cysteine was also excluded from binder sequences. In its default configuration, BindCraft’s designs frequently drifted away from the intended hotspot toward off-target surface locations. We addressed this by increasing the value of BindCraft’s *weights_con_inter* parameter to 2.0 to increase the emphasis placed on contacts at the intended interface during design. We also set *inter_contact_distance* to 8–10 Å (from a default value of 20 Å), and *inter_contact_number* to the number of amino acids listed in the hotspot field (or one less). We found that these changes substantially improved on-target hotspot fidelity of the BindCraft design pipeline (data not shown).

**Table 1.** BindCraft parameter adjustments used in this study to improve on-target hotspot fidelity.

| BindCraft parameter | Default value | Value used in this study |
| --- | --- | --- |
| weights_con_inter | 1.0 | 2.0 |
| inter_contact_distance ($\text{\AA}$) | 20 | 8–10 |
| inter_contact_number | 2 | set to number of amino acids listed in the hotspot field (or one fewer) |

### 2.5 Additional scoring of designed binders with AlphaFold3

Both pipelines include an AF2 module that evaluates binders for independent folding and interaction with the target protein. Since AF3 is newer and more advanced and it would be very challenging for us to replace AF2 with AF3 in either pipeline, we submitted a 295-residue subsequence of 4U40 (given in Supplementary Sequence 1) and each candidate binder as pairs to AFS, which runs AlphaFold3. Although DeepMind has released an AF3 inference pipeline and made model weights available on request, integrating AF3 into either pipeline is impractical due to the added infrastructure required by AF3, so AFS was used instead for evaluation. Each candidate’s binder-target complex was submitted as an AF3 job, and the resulting interface predicted template modeling score (ipTM), predicted template modeling score (pTM), other output parameters, and the predicted aligned error (PAE) plot, were recorded. After the initial set of 15 candidates was evaluated, we conducted a follow-up screen in which five additional candidates with high pipeline ipTM scores were identified from the BindCraft output logs and evaluated by AF3, bringing the total number of AF3-tested candidates to 20 (Table 2). A candidate was considered an “AF3-confirmed” binder only if its AF3-derived ipTM was ≥ 0.80 and its PAE plot displayed a majority of alignment errors under 5 Å (see Results, Section 3.3).

**Table 2.**
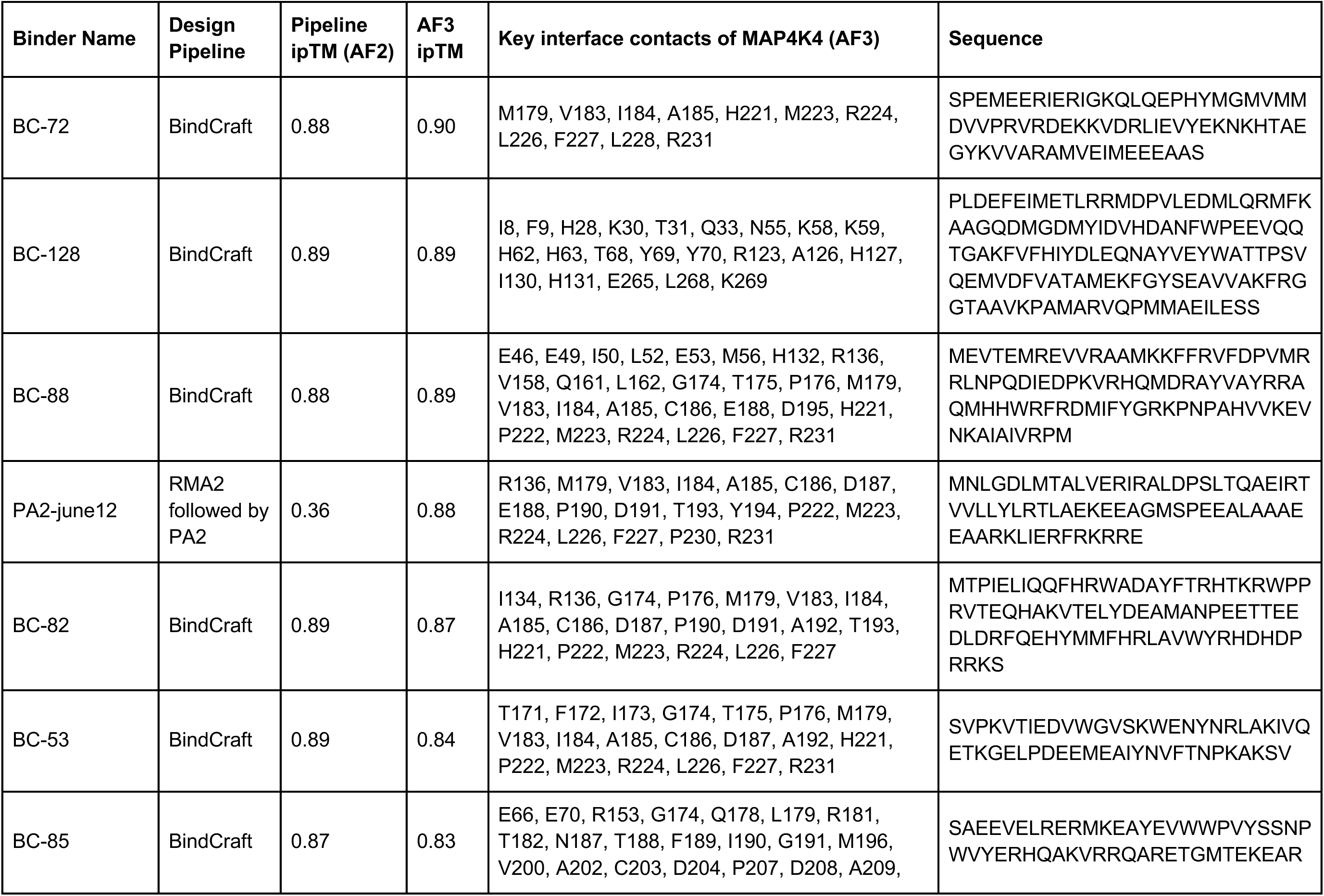
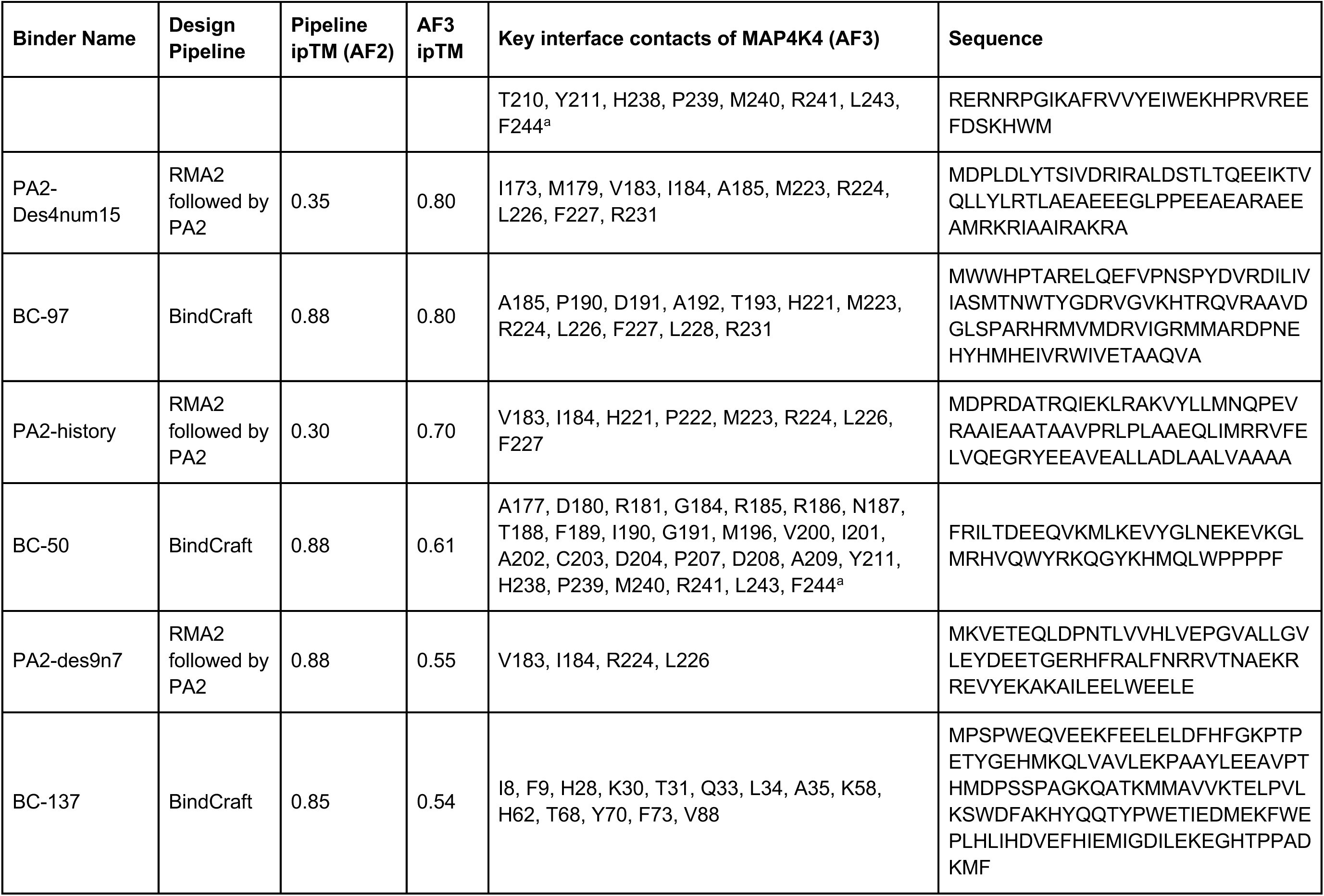
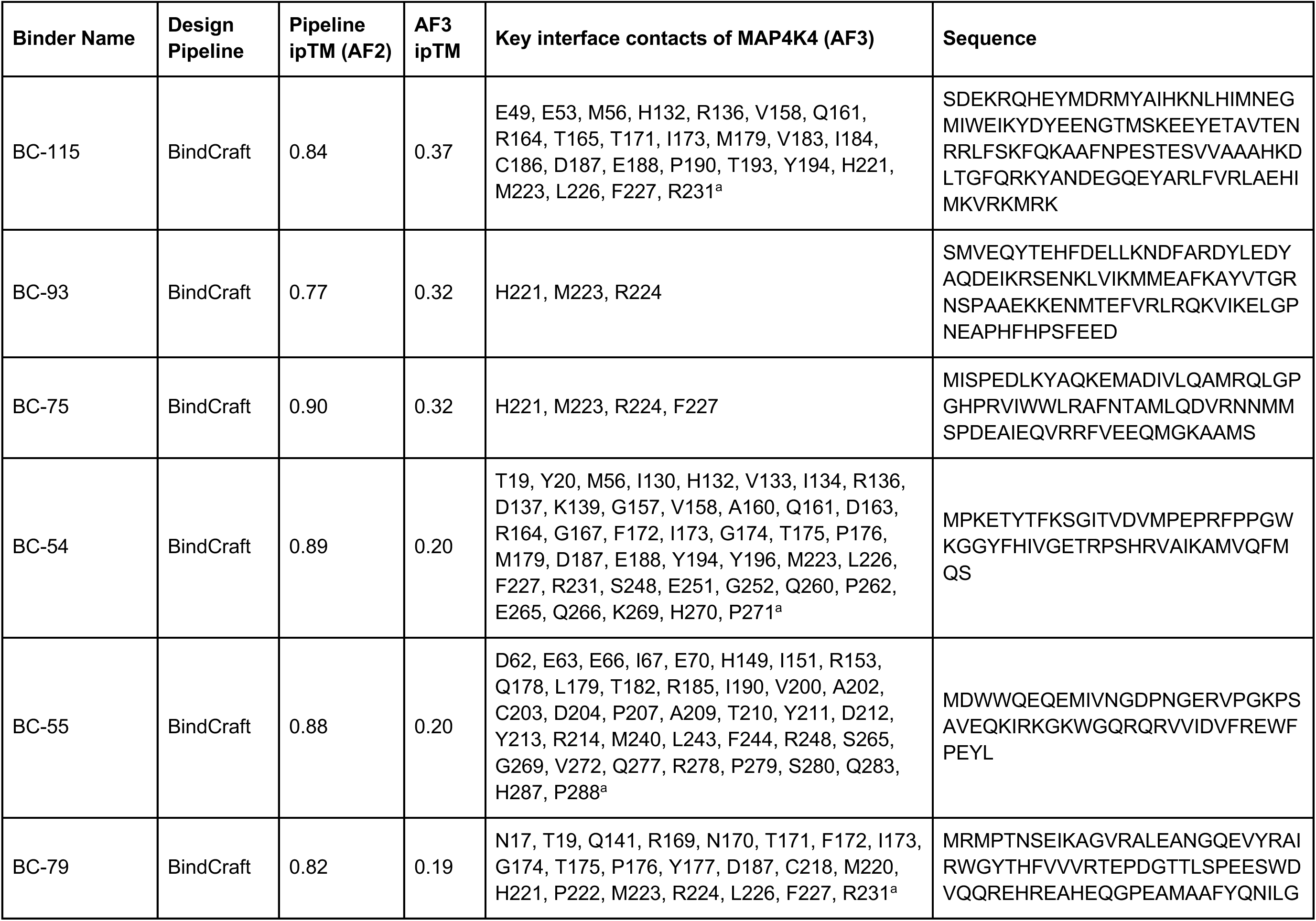
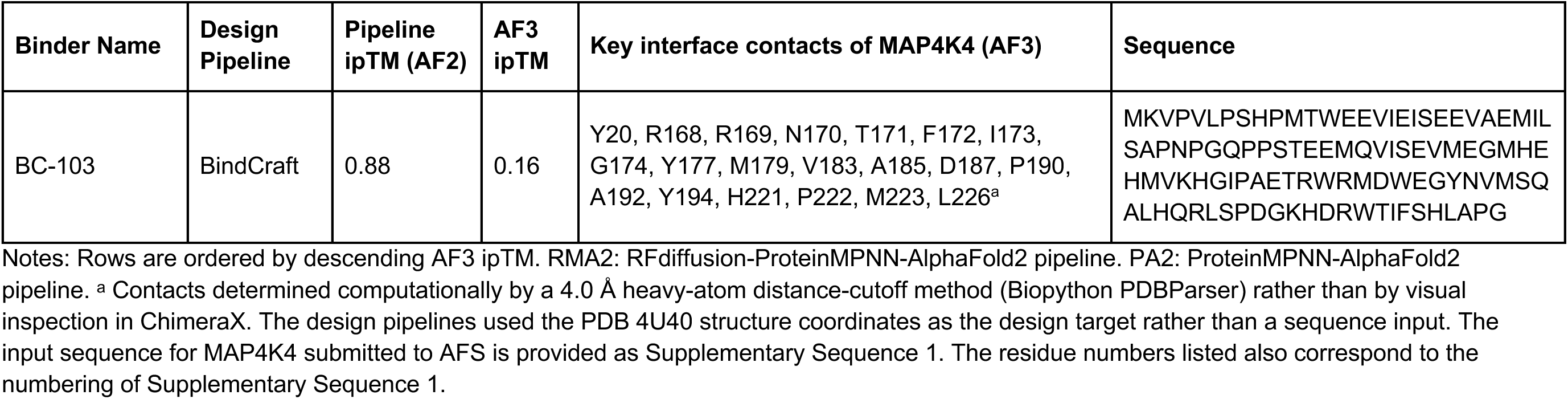
All 20 candidate MAP4K4 binders evaluated in this study.

### 2.6 Structural visualization and sequence novelty

Accepted binder-target complexes were visualized with ChimeraX [17], with MAP4K4 rendered as a tan molecular surface and each binder rendered as a colored ribbon. For each binder, the top-ranked AF3-predicted interface contact residues were mapped onto the structure to confirm that the modeled binding mode matched the intended hotspot; this detailed contact mapping has been completed for the original 15 candidates (Section 3.6). Finally, every designed binder sequence was searched with NCBI BLAST [14,15] to evaluate sequence novelty and rule out rediscovery of known, naturally occurring sequence motifs. The protein BLAST settings were: database: non-redundant (NR) protein database; E-value cutoff threshold: 10; seed word size: 5; matrix: BLOSUM62.

### 2.7 Molecular dynamics simulations

To assess whether the AF3-predicted binding poses of representative candidates remained stable in an explicit-solvent environment, five binder-MAP4K4 complexes spanning the range of AF3-assessed ipTM scores (BC-72, BC-128, BC-88, PA2-june12, and PA2-history) were simulated with GROMACS 2024.3 [19] using the CHARMM36m force field (July 2022 port) [20] and TIP3P water. Each complex model output by AF3 was placed in a cubic box with a 1.0-nm solute-wall padding, solvated explicitly, and neutralized with 0.15 M NaCl. Systems were energy-minimized by steepest descent and equilibrated for 100 ps under an NVT ensemble (V-rescale thermostat, 300 K) followed by 100 ps under an NPT ensemble (C-rescale barostat, 1.0 bar). All bonds to hydrogen were constrained (LINCS) and long-range electrostatics were treated with particle-mesh Ewald (real-space cutoff of 1.0 nm). Production dynamics were run for 100 ns per system (2-fs time step, coordinates saved every 10 ps). Trajectories were analyzed with the GROMACS rms, gyrate, hbond, and rmsf tools to obtain, respectively, whole-complex backbone root-mean-square deviation (RMSD), radius of gyration (R_g_), interchain hydrogen-bond count, and per-residue root-mean-square fluctuation (RMSF), all computed after least-squares fitting to the complex backbone.

### 2.8 AF3 negative-control specificity screening

To evaluate whether the top AF3-scoring binders showed evidence of nonspecific or promiscuous interaction rather than MAP4K4-specific binding, the nine candidates with AF3 ipTM ≥ 0.80 against MAP4K4 (BC-72, BC-128, BC-88, PA2-june12, BC-82, BC-53, BC-85, PA2-Des4num15, and BC-97; see Table 2) were submitted to AFS along with an unrelated human protein kinase negative control, the CDK2 kinase domain (PDB ID: 1HCL, chain A, 298 residues), using the same settings applied to the original MAP4K4 screen. CDK2 was selected as a negative-control target because it is a structurally well-characterized Ser/Thr kinase [21] with no known functional relationship to MAP4K4 and played no role in the hotspot selection used to design these binders. Each binder-CDK2 pair was submitted as an independent AFS job, and output scores and plots were recorded for each of the nine resulting predictions.

## 3. Results

### 3.1 Pipeline yield and AF3 confidence distribution

Across thousands of candidate sequences generated by the two design routes, 20 of the most promising candidates were evaluated by AF3 and are disclosed in Table 2. Fifteen variants emerged from the initial design effort, and 5 additional high-scoring candidates by AF2 were identified in a follow-up search through the design pipeline results (Section 3.6). AF3 ipTM scores for these 20 candidates ranged from 0.16 to 0.90. With MAP4K4 as the target, five candidates (BC-72, BC-128, BC-88, PA2-june12, and BC-82) were scored by AF3 as having ipTM values ≥ 0.87, and four further candidates (ipTM values in parentheses): BC-53 (0.84), BC-85 (0.83), PA2-Des4num15 (0.80), and BC-97 (0.80), extended the high-confidence tier (AF3 ipTM ≥ 0.80) to nine designed binders (Table 2).

### 3.2 Pipeline design and evaluation workflow

Figure 1 summarizes the full workflow used to generate these binders: automated hotspot selection on MAP4K4, design via the RMA2/PA2 or BindCraft pipeline, AF3 evaluation, final structural visualization, and submission to NCBI BLAST for a sequence novelty check.

### 3.3 Pipeline AF2 scores do not reliably predict AF3 values

Pipeline-stage parameter values determined by AF2 frequently diverged from the AF3-derived results for the same output sequences. BC-79 scored a pipeline ipTM of 0.82 but collapsed to an AF3 ipTM of only 0.19; BC-93 similarly fell from a pipeline AF2 ipTM of 0.77 to an AF3 ipTM of 0.32 (Table 2). In contrast, BC-128 scored 0.89 at the pipeline stage and held at 0.89 with AF3. Across the full candidate set, several candidates with strong pipeline ipTM values (BC-75, 0.90; PA2-des9n7, 0.88) scored poorly with AF3 (0.32 and 0.55, respectively), while PA2-june12 and PA2-Des4num15, both with modest pipeline ipTM scores (0.36 and 0.35), scored much higher with AF3 (0.88 and 0.80). Across all 20 candidates, AF3 ipTM was lower than the pipeline AF2 ipTM for 14 (70%), higher for 5 (25%), and equal for 1 (5%), with essentially no positive correlation between the two scores (Pearson r = −0.25; Figure 5); this is consistent with AF3 generally functioning as a harder grader than the in-pipeline AF2 module (Section 4).

### 3.4 Structural modeling of top candidates

Figure 2 shows ChimeraX renderings of four binders bound to MAP4K4, including each binder’s top-ranked interface contact residues as determined by AF3. BC-128 and PA2-june12 engage distinct, non-overlapping surfaces of MAP4K4 (contacts I8/F9/H28/K30 versus R136/M179/V183/I184, respectively), on opposite faces of the target (Figure 2E), making this pair a candidate for a two-site sandwich assay.

**Figure 2.**
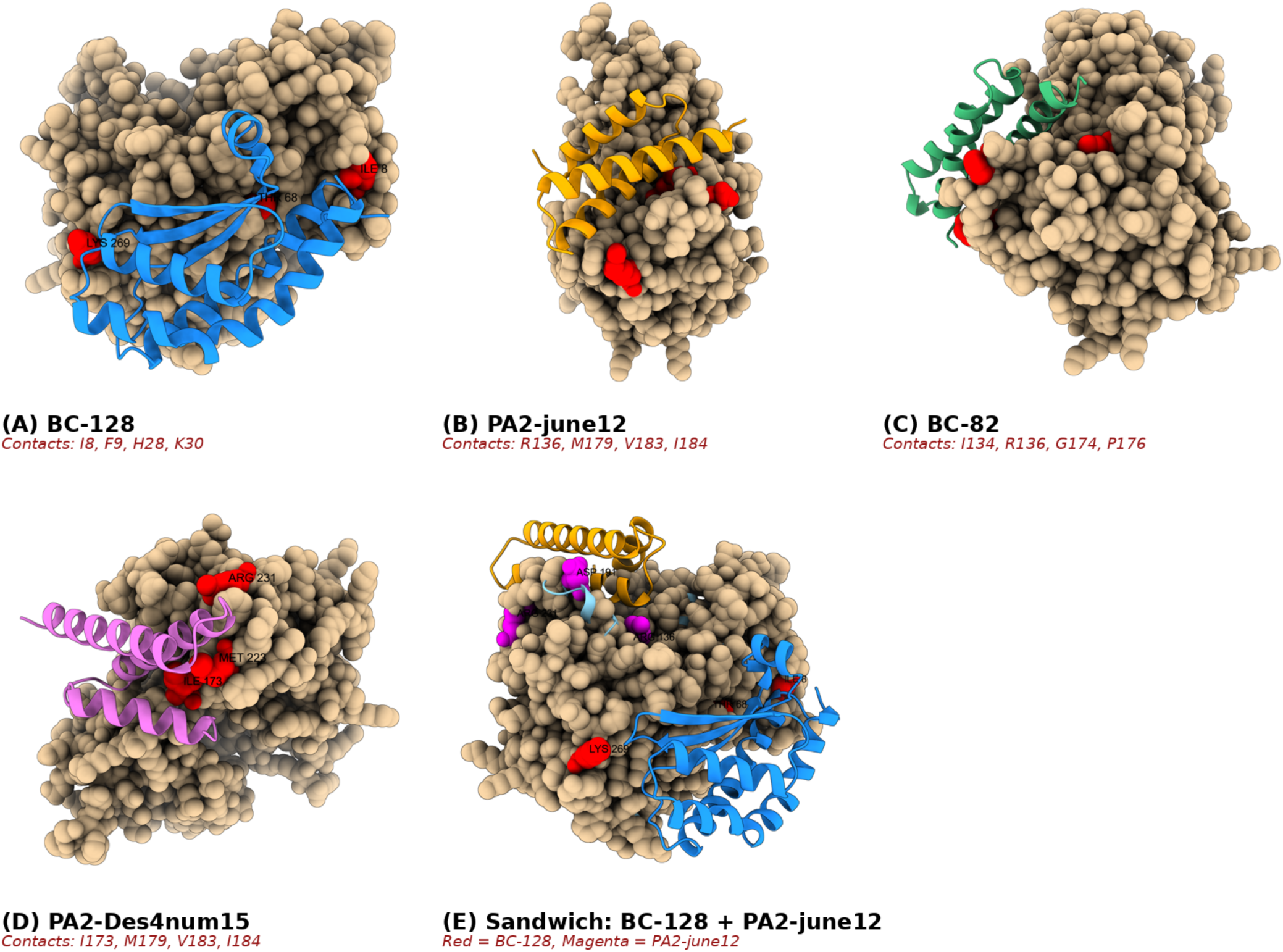
ChimeraX renderings of four MAP4K4 binders. MAP4K4 is shown as a tan molecular surface; each designed binder is shown as a colored ribbon. Red/magenta spheres mark each binder’s top-ranked interface contact residues as determined by AF3. (A) BC-128. (B) PA2-june12. (C) BC-82. (D) PA2-Des4num15. All interface contact residues are listed in Table 2. (E) Composite view of BC-128 (red) and PA2-june12 (magenta) bound simultaneously to non-overlapping faces of MAP4K4, illustrating their potential as a two-site sandwich-assay pair.

### 3.5 Confidence assessment via predicted aligned error

To further assess the reliability of the six highest-confidence candidates from the initial screen, we examined the Predicted Aligned Error (PAE) plots returned by AFS for each independent evaluation (Figure 3). Low predicted error in the off-diagonal rectangular blocks corresponding to the interface between the binder and the target, for each of the six top candidates, indicates that the relative position and orientation of binder and target were modeled with high confidence, consistent with the corresponding high ipTM and pTM scores (Table 2).

**Figure 3.**
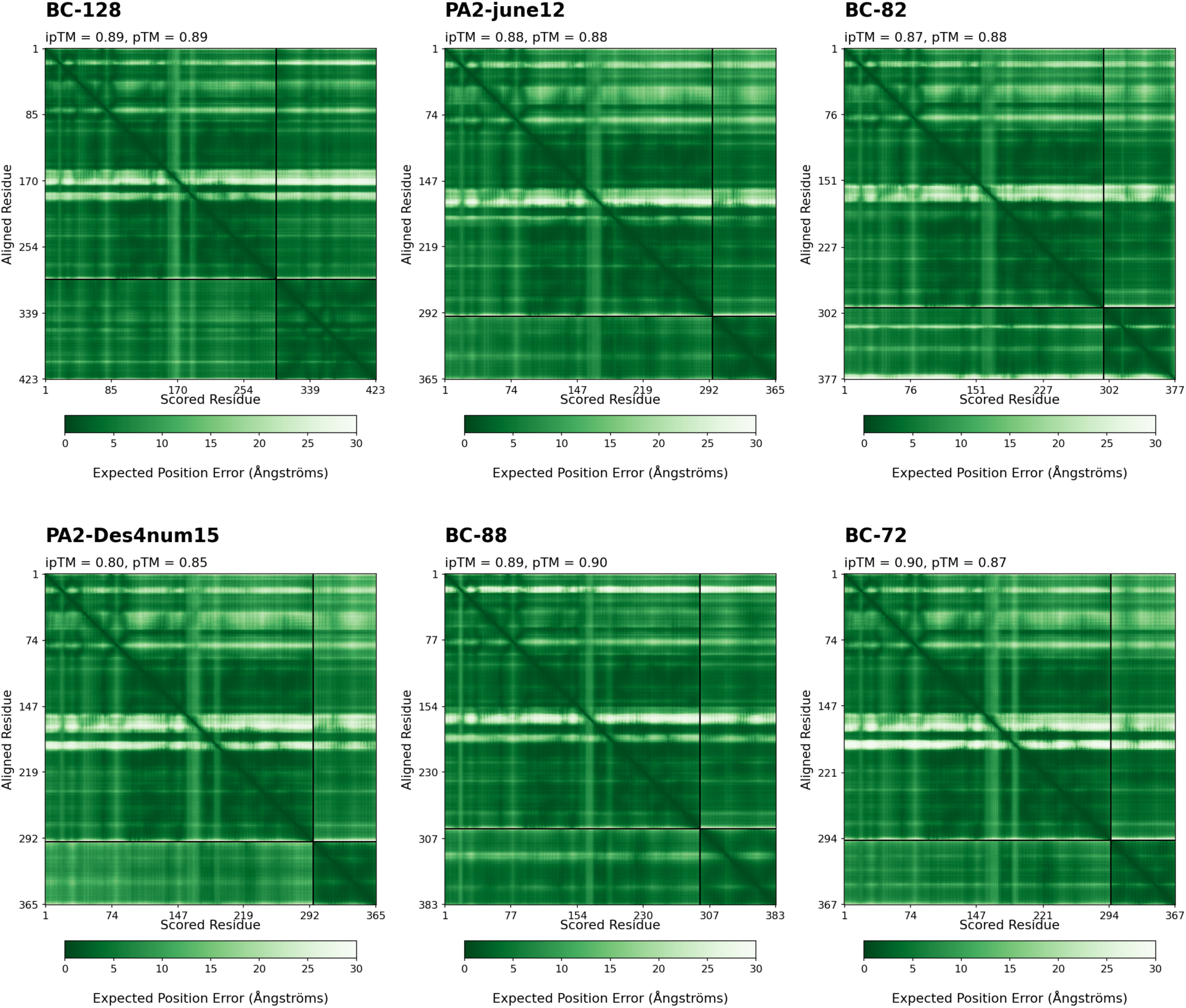
Predicted Aligned Error (PAE) plots from AlphaFold Server (AF3) evaluation of six high-confidence MAP4K4 binders identified in the initial screen. Each heatmap plots expected position error (Å, 0–30 Å scale) for every pair of residues in the modeled complex; low error in the off-diagonal blocks corresponding to the binder-target interface (separated by the chain-boundary divider line) indicates confident, well-defined relative positioning between binder and target chains. Panels are labeled with binder identity and AF3 ipTM/pTM scores.

### 3.6 Extended candidate search identifies one additional high-confidence binder

To test whether additional strong binders remained, we mined the BindCraft output logs for the five candidates with the highest self-reported pipeline ipTM that had not yet been submitted to AF3. This effort yielded BC-54 (AF2 ipTM of 0.89), BC-103 (0.88), BC-55 (0.88), BC-50 (0.88), and BC-85 (0.87). We submitted all five for scoring by AF3. Consistent with the pattern described in Section 3.3, four of the five scored poorly: BC-54 (AF3 ipTM 0.20, pTM 0.75), BC-103 (AF3 ipTM 0.16, pTM 0.67), BC-55 (AF3 ipTM 0.20, pTM 0.71), and BC-50 (AF3 ipTM 0.61, pTM 0.77) and fell well short of their pipeline scores. BC-85, however, scored highly (AF3 ipTM 0.83, pTM 0.82), joining the high-confidence tier of AF3-confirmed binders.

### 3.7 Sequence novelty

NCBI BLASTp searches [14,15] of all 20 designed binder sequences in Table 2 against its non-redundant protein sequences database returned no matches to known proteins for ten of the binders and low-significance matches for the other ten (minimum E > 0.6, Table S2). The matching sequences were all of microbial origin, with no connection to MAP4K4. These results are consistent with the designed binders being truly novel protein sequences rather than rediscoveries of known proteins or motifs, or even chimeras or other amalgamations of known proteins.

### 3.8 Stability assessment of top candidate complexes by molecular dynamics

Five AF3-evaluated complexes of MAP4K4 with designed binder candidates (BC-72, BC-128, BC-88, PA2-june12, and PA2-history) were carried forward into 100-ns explicit-solvent GROMACS simulations to test whether their AF3-predicted binding poses persisted beyond a single static structural prediction (Figure 4). Across all five systems, at least one interchain hydrogen bond was present in essentially every sampled frame (Figure 4C, ≥99.9% of frames for all five systems), indicating that the designed binding interface was never fully released over the full 100-ns trajectory. Whole-complex backbone root-mean-square deviation (RMSD), computed after least-squares fitting to the complex backbone, varied considerably between systems and over time (Figure 4A). BC-88 was the most rigidly engaged complex by every metric. It possessed the lowest whole-complex RMSD of the five systems (0.27 ± 0.57 nm over the full trajectory, with only brief excursions to a maximum of 3.6 nm) and held the highest sustained interchain hydrogen-bond count (11.1 ± 2.2 bonds, full trajectory). BC-128 became progressively less stable over the run (backbone RMSD 1.37 ± 1.62 nm over the full trajectory, rising to 2.56 ± 1.54 nm over the second half of the simulated period, with a 5.47-nm maximum, the largest of the five systems). The RMSD of BC-72 remained elevated throughout (2.38 ± 1.43 nm full trajectory; 2.06 ± 1.34 nm second half). Variant PA2-june12, despite substantial early-trajectory reorientation, settled into a markedly lower-RMSD pose over the second half of the simulation (0.22 ± 0.39 nm for 50–100 ns versus 1.22 ± 1.62 nm over the full trajectory), suggesting the complex reached a new, comparatively stable orientation after an initial adjustment period. PA2-history, the lowest-ipTM system subjected to MD (AF3 ipTM 0.70), exhibited intermediate RMSD values throughout (0.47 ± 0.85 nm full trajectory; 0.74 ± 1.14 nm in the second half).

**Figure 4.**
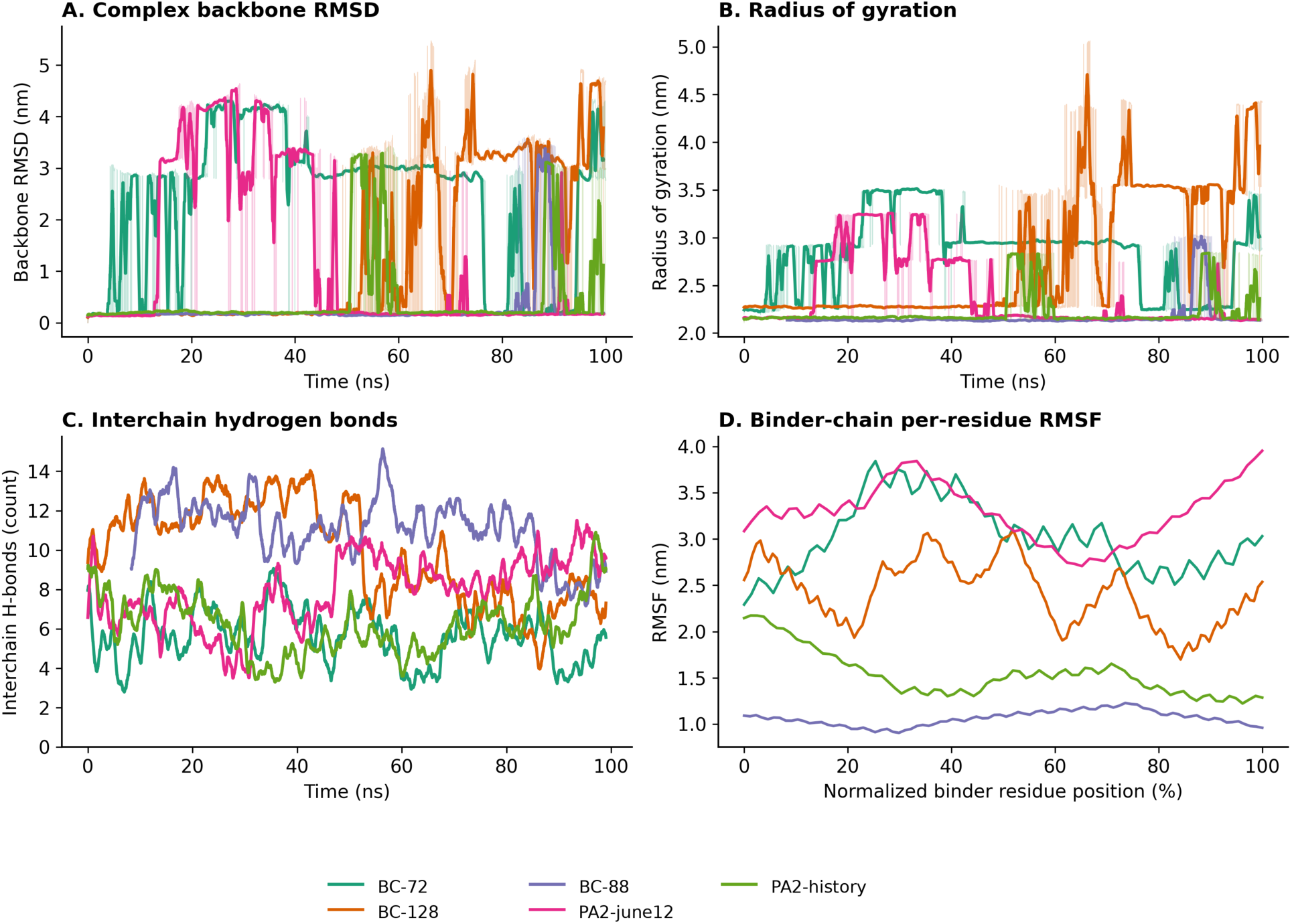
Molecular dynamics stability plots for five top-scoring MAP4K4-binder complexes over 100 ns of explicit-solvent GROMACS simulation. (A) Whole-complex backbone RMSD versus time (light traces: raw 10-ps resolution data; heavy traces: 0.51-ns moving average). (B) Radius of gyration versus time (same smoothing convention as panel A). (C) Interchain hydrogen-bond count versus time (1.01 ns moving average). (D) Per-residue RMSF of the binder chain only, plotted against normalized residue position along each binder’s sequence, with each binder’s chain length given in the legend. All panels computed after least-squares fitting to the complex backbone; see Section 2.7 for simulation parameters and Section 3.8 for discussion.

Final-frame backbone RMSD values reinforced this picture: BC-128 ended the 100 ns trajectory at 4.68 nm, close to its 5.47 nm maximum, while BC-88 and PA2-june12 both ended near their respective full-trajectory minima (0.18 nm and 0.16 nm). PA2-history was an exception to this otherwise consistent picture: despite a low full-trajectory mean RMSD (0.47 ± 0.85 nm), it ended the simulation at an elevated 2.97 nm, indicating a late-stage excursion whose long-term stability was not resolved within the simulated 100 ns window.

Per-residue root-mean-square fluctuation (RMSF, Figure 4D), computed for each chain after the same whole-complex fit, displayed an analogous pattern to RMSD: elevated RMSF was driven predominantly by motion of the small binder chain relative to the much larger ∼295-residue MAP4K4 protein, rather than by internal unfolding of either chain. Mean MAP4K4 RMSF ranged from 0.37 nm (BC-88) to 1.51 nm (BC-72), while mean binder RMSF ranged from 1.06 nm (BC-88) to 3.30 nm (PA2-june12) (Figure 4D).

Radius of gyration (R_g_, Figure 4B) stayed within a bounded range for every system rather than diverging over time, consistent with reorientation of an intact complex rather than dissociation or unfolding. Radius of gyration is a measure of overall complex compactness: full-trajectory R_g_ ranged from 2.16 ± 0.16 nm (BC-88) to 2.84 ± 0.41 nm (BC-72), and stayed close to its full-trajectory mean in the second half for four of the five systems (shift ≤ 0.23 nm); the exception was BC-128, whose R_g_ rose from 2.77 ± 0.69 nm (full trajectory) to 3.27 ± 0.68 nm (second half), corroborating the progressive destabilization already indicated by its rising RMSD.

Interchain hydrogen-bond counts diverged between the first and second half of the trajectory in the same direction as the RMSD-based stability ranking: BC-128 lost contacts over time (9.9 ± 2.9 bonds full trajectory, falling to 7.9 ± 2.2 in the second half), while PA2-june12 gained them (7.9 ± 2.3 bonds full trajectory, rising to 9.2 ± 1.7 in the second half), consistent with its settling into the lower-RMSD pose noted above.

Taken together, these simulations indicate that interchain contact was retained for all five designs over 100 ns of explicit-solvent dynamics, with BC-88 standing out as the most conformationally rigid complex and, therefore, the strongest MD-supported candidate among the five systems tested.

### 3.9 Binder specificity: negative-control screening against CDK2

To test whether any of the nine candidates with the highest AF3 ipTM scores and favorable PAE plots (AF3 ipTM ≥ 0.80; Table 2) interact with MAP4K4 in a non-specific or promiscuous manner, each was independently evaluated for binding to a negative-control target, CDK2 (Section 2.8). All nine binder-CDK2 predictions returned substantially lower confidence parameter values than the corresponding binder-MAP4K4 predictions (results listed in Table 3).

**Table 3.** AF3 negative-control specificity screen: ipTM against MAP4K4 versus CDK2 for the nine highest-confidence binders.

| Binder Name | AF3 ipTM vs. MAP4K4 | AF3 ipTM vs. CDK2 (negative control) |
| --- | --- | --- |
| BC-72 | 0.90 | 0.11 |
| BC-128 | 0.89 | 0.13 |
| BC-88 | 0.89 | 0.15 |
| PA2-june12 | 0.88 | 0.24 |
| BC-82 | 0.87 | 0.19 |
| BC-53 | 0.84 | 0.12 |
| BC-85 | 0.83 | 0.13 |
| PA2-Des4num15 | 0.80 | 0.20 |
| BC-97 | 0.80 | 0.10 |

Notes: CDK2 (PDB ID: 1HCL, chain A) served as the negative-control target (Section 2.8). ipTM against MAP4K4 exceeded ipTM against CDK2 for all nine binders (Wilcoxon signed-rank test, paired by binder: $W = 0$ , $n = 9$ , $p = 0.0039$ ; paired t-test: $t(8) = 35.9$ , $p < 0.0001$ ).

Across all nine candidates, CDK2 ipTM ranged from 0.10 to 0.24 (mean 0.15), an approximately 4- to 8-fold reduction from the 0.80–0.90 AF3 ipTM range these same binders achieved against MAP4K4 (Table 3), with ipTM against MAP4K4 substantially exceeding ipTM against CDK2 for every one of the nine binders tested; this difference was statistically significant (see Table 3 note). The large reduction in ipTM against an unrelated kinase across all nine candidates is consistent with MAP4K4-specific rather than nonspecific or promiscuous binding.

### 3.10 Predicted biophysical properties of top candidates

Beyond the structural and interface metrics reported above, the nine AF3-confirmed candidates (Table 2) were evaluated for physicochemical properties relevant to production and formulation. Standard ProtParam descriptors, computed from each binder sequence, are summarized in Supplementary Table S1. Instability index values exceeded the conventional threshold of 40, below which a protein is classified as unstable by the ExPASy criterion, for eight of the nine candidates, ranging from 43.9 (BC-128) to 106.1 (PA2-june12); only BC-53 fell below this threshold, at 22.6. This index is calibrated on a reference set of natural globular proteins, and small, highly charged, largely helical designs such as these routinely fall outside that calibration space regardless of their actual folding behavior, so we treat this as a minor flag.

Toxicity was assessed with ToxinPred2 [22] (hybrid model, default threshold 0.6; Supplementary Table S1). Two of nine candidates were classified as toxic: PA2-june12 (score 0.67) and PA2-Des4num15 (score 0.76); the remaining seven, including all candidates from the BindCraft pipeline, were classified as non-toxic. ToxinPred2 is trained primarily on natural venom and bacterial toxin sequences. These two designs’ small size, high net charge, and predominantly helical content compositionally overlap several natural peptide toxin classes despite having no evolutionary relationship to them. As such, it is again likely that these flags were raised due to calibration mismatches. As with all computational predictions, however wet lab assays would be needed to confirm toxicity assignments.

Allergenicity was assessed with AlgPred 2.0 [23] (hybrid model, default threshold 0.3; Supplementary Table S1). One candidate, BC-88, was classified as an allergen (hybrid score 0.69). Decomposing this call shows it was driven almost entirely by a MERCI motif-search score of 0.5 against known allergen sequence motifs; the underlying AlgPred 2.0 machine-learning score for BC-88 alone was 0.19 (Table S1), below the classification threshold [23].

Considered by pipeline of origin, both toxicity flags arose from candidates produced by the RFdiffusion-ProteinMPNN-AlphaFold2 (RMA2) route (PA2-june12, PA2-Des4num15), while none of the seven BindCraft-derived candidates were flagged. Given the small number of candidates per pipeline in this set (seven BindCraft, two RMA2-derived), this is simply an observation rather than a confirmed difference in the two pipelines.

Finally, these flags are not confined to lower-ranked candidates. BC-88, which carries the allergenicity flag, is the third-ranked candidate by AF3 ipTM (0.89); PA2-june12, which carries a toxicity flag, is ranked fourth (ipTM 0.88). Two of the four highest-confidence candidates by AF3 ipTM therefore each carry one predicted flag, underscoring that AF3 confidence and the biophysical screens applied here assess different properties and should be considered jointly rather than in place of one another.

## 4. Discussion

A noteworthy methodological finding of this study is that AF3, a newer and generally more conservative evaluator than AF2 [24], the in-pipeline module for the design tools used in this work, output ipTM values that frequently disagreed with AF2 ipTM scores (Figure 5). Several candidates with strong pipeline-stage AF2 ipTM (e.g., BC-75 and BC-79, both ≥ 0.82; and, in the follow-up screen, BC-54, BC-103, and BC-55, all ≥ 0.88) failed to score highly by AF3, while others with weak pipeline scores (PA2-june12, PA2-Des4num15, and PA2-history; all < 0.4) scored much higher with AF3. The former suggests that pipeline-stage AF2 scores can reflect “overconfidence” in or “gaming” of a pipeline’s evaluation method. This pattern echoes the broader phenomenon in which a metric used as an internal optimization target tends to become a less reliable indicator of the underlying quality it was meant to track [25]. An iterative, largely self-directed pipeline like BindCraft may be especially prone to generating proteins with sequence features that “artificially” inflate its own evaluation tool’s confidence score. The three proteins with much higher ipTM values from AF3 vs. from AF2 are more curious and also demonstrate the value of submitting as many *de novo* binder candidates as possible to additional evaluation tools. Though it may be possible to amend the RFdiffusion-ProteinMPNN-AF2 and BindCraft pipelines to run AF3, the newer and more respected tool, instead of AF2 for the binder-target complex evaluation stage, such a change is far from trivial, especially for teams like ours without research funding. We therefore relied on manual submission of binder-target sequence pairs to the AlphaFold Server for our AF3-based evaluations.

**Figure 5.**
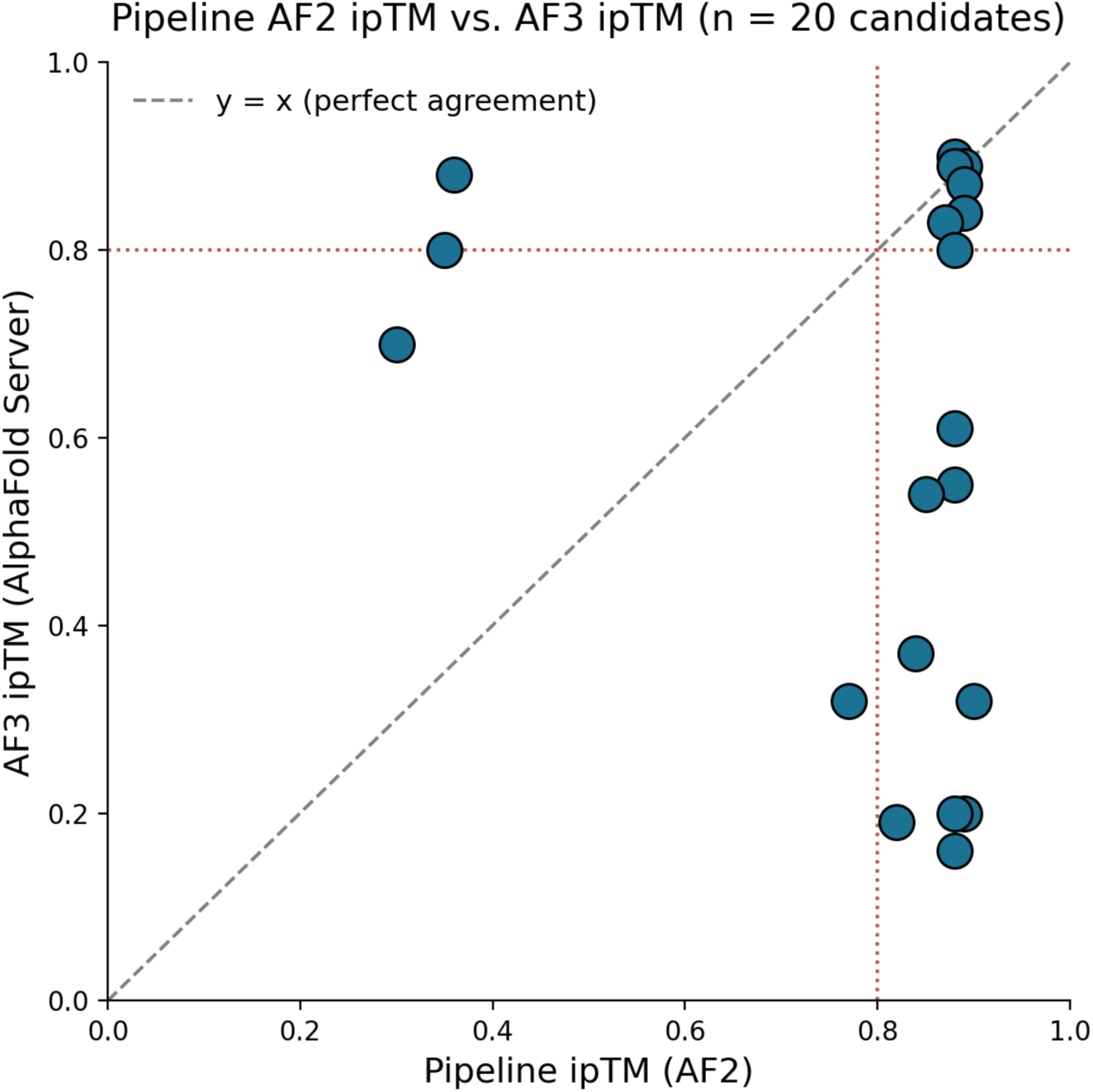
AF3 ipTM versus AF2 ipTM for all 20 evaluated candidates. Scatter plot of pipeline-stage AF2 ipTM versus AF3 ipTM for all 20 candidates in Table 2 (PA2-history was rescored with a standalone single-pair AF2-multimer evaluation; Section 3.3). Dotted line at 0.80 marks the AF3-confirmed threshold on each axis; dashed diagonal marks perfect agreement. AF3 ipTM was lower than the pipeline AF2 ipTM for 14 of 20 candidates (70%), higher for 5 (25%), and equal for 1 (5%); Pearson r = −0.25, indicating essentially no positive correlation between the two scores.

Beyond the AF3 check, we added two further computational tests for the highest-confidence candidates. Explicit-solvent MD simulations of five representative complexes (Section 3.8) showed that interchain contact persisted throughout 100 ns for every system tested, and radius of gyration and interchain hydrogen-bonding trends independently corroborated the RMSD-based stability ranking. Two complexes with MAP4K4, BC-128 and PA2-history, showed continued conformational drift that had not resolved by 100 ns, underscoring that a single simulated timescale cannot rule out slower rearrangements. Second, evaluation of the nine highest-confidence binders against a negative control target protein, CDK2 (Section 3.9), produced a consistent, statistically significant drop in ipTM relative to MAP4K4 (Wilcoxon signed-rank test, *p* = 0.0039), supporting that these interactions are specific to MAP4K4, rather than an artifact of generically “sticky” designed surfaces. Neither additional test replaces wet-lab binding and specificity assays, but together with the AF3 evaluation, they comprise provide substantial additional support that candidate target-binding proteins are strong prospects for experimental follow-up.

The follow-up computational screen described in Section 3.6 makes this point especially concrete: of five additional candidates chosen purely because they had the highest pipeline AF2-reported ipTM values, four (BC-54, BC-103, BC-55, and BC-50) scored poorly, with AF3 ipTM values of 0.16–0.61 and only one, BC-85, held up (AF3 ipTM of 0.83). Selecting candidates for AF3 testing by pipeline AF2 score alone would have missed the single strongest such case, variant PA2-june12 (AF2 ipTM of 0.36; AF3 ipTM of 0.88) and would have wasted evaluation resources on the four variants listed above whose AF3 ipTM values were markedly lower than their corresponding AF2 scores. This teaches that broad AF3 screening of many candidates, not only the very highest-scoring designs by AF2, can allow for discovery of quality binder candidates that would otherwise be discarded.

Our finding, though not rigorously documented, that relatively simple model parameter adjustments (tuning BindCraft’s weights_con_inter, inter_contact_distance, and inter_contact_number) meaningfully improved on-target hotspot fidelity is notable because it required no change to the underlying design models, only to how strongly the optimization engine weighted the intended interface. This suggests that one-shot design pipelines like BindCraft, while convenient, may need small amounts of target-specific tuning to reliably anchor designs at a chosen hotspot, and that such tuning can be identified without specialized computational infrastructure. It is valuable to note, however, that BindCraft was significantly more computationally expensive, especially when attempting to generate smaller binders.

That both the RMA2/PA2 and one-shot (BindCraft) design pipelines independently produced binders that achieved high (>0.80) ipTM values from AF3, several of which converge on overlapping surface regions of MAP4K4 (e.g., contacts near residues 179-185 recur in BC-72, PA2-june12, BC-53, and PA2-Des4num15; see Table 2), supports the reproducibility of these hotspots as genuine, potentially targetable epitopes on MAP4K4 rather than artifacts of a single design method.

The set of AF3-confirmed hotspots maps several distinct surface regions of MAP4K4 that have not been previously characterized as functional sites. Contacts clustering around residues 179 to 231 recur across four independently designed binders (BC-72, PA2-june12, BC-53, and PA2-Des4num15), while BC-128 engages a separate cluster near residues 8, 9, 28, 30, and 33, and BC-82 engages a third cluster near residues 134, 136, and 174 to 176. Some of these binder-defined surfaces do not appear to be targeted by existing characterized antibodies (authors’ investigation, data not shown), suggesting the clusters mark regions of the MAP4K4 surface that remain relatively unexplored as engineered-binder targets. A structurally diverse binder panel of this kind could therefore serve as a set of site-specific probes for researchers to map which regions of the MAP4K4 surface participate in its various signaling activities across the JNK, p38 MAPK, and ERK1/2 pathways, thus offering a route toward distinguishing functionally distinct sites on a single, multi-pathway kinase.

This work has several limitations. No binder reported herein has yet been tested in a wet-lab binding assay. Promising AF3 output values are strong computational evidence of protein folding and interaction, but are not a substitute for experimental validation, such as with surface plasmon resonance (SPR), biolayer interferometry (BLI), or ELISA-type assays. The hotspot-scoring weights were tuned specifically for MAP4K4 and have not yet been tested on other targets, so their generalizability is unknown. On the other hand, the BindCraft parameter adjustments used here to promote “adherence” to the specified hotspot have been used with four other target proteins, to beneficial effect (results not shown). Finally, the 20 candidates reported here result from a limited number of hotspot-selection rounds rather than a systematic, full-surface panel, so additional high-confidence binder sites on MAP4K4 likely remain undiscovered. Predicted biophysical properties (Section 3.10) should also be interpreted with the same caution as the instability index, as several of the underlying tools are calibrated on natural proteins, which are products of evolution, and may not generalize cleanly to small *de novo* designs, which are not.

## 5. Conclusions

Taken together, the binder-defined hotspots also sketch an early, data-driven map of the MAP4K4 surface that may point toward previously uncharacterized biologically active sites. Future work should prioritize wet-lab validation and binding affinity determination of the top candidates, assessment of a sandwich assay comprising the BC-128/PA2-june12 pair, ChimeraX visual confirmation of the computationally assigned contacts for BC-85 and its follow-up-screen counterparts (Table 2, note a), expansion of hotspot coverage to a full-surface panel, systematic comparison of the binder-defined hotspot clusters against known regulatory and protein-interaction surfaces to determine which correspond to distinct biologically active sites on MAP4K4, and, if justified, cell-based assays testing whether these binders modulate MAP4K4-dependent JNK, p38, and ERK1/2 signaling.

Despite its limitations, this study demonstrates that *de novo* protein-binder design against a therapeutically relevant human kinase can be performed using only free, publicly available computational tools, without research grants or access to a well-resourced laboratory. By pairing these design pipelines with binding interaction assessment by AlphaFold3 and molecular dynamics simulations, this research provides additional support for predicted binder folding and target-binding functionality.

## Author Contributions

A.J.: conceptualization, methodology, custom analysis and scoring-tool code development, formal analysis, investigation, data curation, visualization, writing, composing original draft. A.T.: conceptualization, supervision, instruction, methodology, writing, review and editing.

## Acknowledgments

The authors thank Sergey Ovchinnikov and Martin Pacesa for creating and posting the RFdiffusion and BindCraft pipelines, respectively, for free use in Google Colaboratory.

## References

1. Singh, S.K., Roy, R., Kumar, S., Srivastava, P., Jha, S., Rana, B., Rana, A. Molecular Insights of MAP4K4 Signaling in Inflammatory and Malignant Diseases. Cancers 15, 2272 (2023).

2. Bouzakri, K., Zierath, J.R. MAP4K4 Gene Silencing in Human Skeletal Muscle Prevents Tumor Necrosis Factor-α-Induced Insulin Resistance. J. Biol. Chem. 282, 7783–7789 (2007).

3. Aouadi, M., Tesz, G.J., Nicoloro, S.M., Wang, M., Chouinard, M., Soto, E., Ostroff, G.R., Czech, M.P. Orally Delivered siRNA Targeting Macrophage Map4k4 Suppresses Systemic Inflammation. Nature 458, 1180–1184 (2009).

4. Chuang, H.-C., Sheu, W.H.-H., Lin, Y.-T., Tsai, C.-Y., Yang, C.-Y., Cheng, Y.-J., Huang, P.-Y., Li, J.-P., Chiu, L.-L., Wang, X., Xie, M., Schneider, M.D., Tan, T.-H. HGK/MAP4K4 Deficiency Induces TRAF2 Stabilization and Th17 Differentiation Leading to Insulin Resistance. Nat. Commun. 5, 4602 (2014).

5. Fiedler, L.R., Chapman, K., Xie, M., Maifoshie, E., Jenkins, M., Golforoush, P.A., Bellahcene, M., Noseda, M., Faust, D., Jarvis, A., Newton, G., Paiva, M.A., Harada, M., Stuckey, D.J., Song, W., Habib, J., Narasimham, P., Aqil, R., Sanmugalingam, D., Yan, R., Pavanello, L., Sano, M., Wang, S.C., Sampson, R.D., Kanayaganam, S., Taffet, G.E., Michael, L.H., Entman, M.L., Tan, T.-H., Harding, S.E., Low, C.M.R., Tralau-Stewart, C., Perrior, T., Schneider, M.D. MAP4K4 Inhibition Promotes Survival of Human Stem Cell-Derived Cardiomyocytes and Reduces Infarct Size In Vivo. Cell Stem Cell 24, 579–591 (2019).

6. Kahn, R.A., Virk, H., Laflamme, C., Houston, D.W., Polinski, N.K., Meijers, R., Levey, A.I., Saper, C.B., Errington, T.M., Turn, R.E., Bandrowski, A., Trimmer, J.S., Rego, M., Freedman, L.P., Ferrara, F., Bradbury, A.R.M., Cable, H., Longworth, S. Antibody characterization is critical to enhance reproducibility in biomedical research. eLife 13, e100211 (2024).

7. Watson, J.L., Juergens, D., Bennett, N.R., Trippe, B.L., Yim, J., Eisenach, H.E., Ahern, W., Borst, A.J., Ragotte, R.J., Milles, L.F., Wicky, B.I.M., Hanikel, N., Pellock, S.J., Courbet, A., Sheffler, W., Wang, J., Venkatesh, P., Sappington, I., Vázquez Torres, S., Lauko, A., De Bortoli, V., Mathieu, E., Ovchinnikov, S., Barzilay, R., Jaakkola, T.S., DiMaio, F., Baek, M., Baker, D. De Novo Design of Protein Structure and Function with RFdiffusion. Nature 620, 1089–1100 (2023).

8. Dauparas, J., Anishchenko, I., Bennett, N., Bai, H., Ragotte, R.J., Milles, L.F., Wicky, B.I.M., Courbet, A., de Haas, R.J., Bethel, N., Leung, P.J.Y., Huddy, T.F., Pellock, S., Tischer, D., Chan, F., Koepnick, B., Nguyen, H., Kang, A., Sankaran, B., Bera, A.K., King, N.P., Baker, D. Robust Deep Learning-Based Protein Sequence Design Using ProteinMPNN. Science 378, 49–56 (2022).

9. Jumper, J., Evans, R., Pritzel, A., Green, T., Figurnov, M., Ronneberger, O., Tunyasuvunakool, K., Bates, R., Žídek, A., Potapenko, A., Bridgland, A., Meyer, C., Kohl, S.A.A., Ballard, A.J., Cowie, A., Romera-Paredes, B., Nikolov, S., Jain, R., Adler, J., Back, T., Petersen, S., Reiman, D., Clancy, E., Zielinski, M., Steinegger, M., Pacholska, M., Berghammer, T., Bodenstein, S., Silver, D., Vinyals, O., Senior, A.W., Kavukcuoglu, K., Kohli, P., Hassabis, D. Highly Accurate Protein Structure Prediction with AlphaFold. Nature 596, 583–589 (2021).

10. Pacesa, M., Nickel, L., Schellhaas, C., Schmidt, J., Pyatova, E., Kissling, L., Barendse, P., Choudhury, J., Kapoor, S., Alcaraz-Serna, A., Cho, Y., Ghamary, K.H., Vinué, L., Yachnin, B.J., Wollacott, A.M., Buckley, S., Westphal, A.H., Lindhoud, S., Georgeon, S., Goverde, C.A., Hatzopoulos, G.N., Gönczy, P., Muller, Y.D., Schwank, G., Swarts, D.C., Vecchio, A.J., Schneider, B.L., Ovchinnikov, S., Correia, B.E. BindCraft: One-Shot Design of Functional Protein Binders. bioRxiv (2024). https://github.com/martinpacesa/BindCraft

11. Abramson, J., Adler, J., Dunger, J., Evans, R., Green, T., Pritzel, A., Ronneberger, O., Willmore, L., Ballard, A.J., Bambrick, J., Bodenstein, S.W., Evans, D.A., Hung, C.-C., O’Neill, M., Reiman, D., Tunyasuvunakool, K., Wu, Z., Žemgulytė, A., Arvaniti, E., Beattie, C., Bertolli, O., Bridgland, A., Cherepanov, A., Congreve, M., Cowen-Rivers, A.I., Cowie, A., Figurnov, M., Fuchs, F.B., Gladman, H., Jain, R., Khan, Y.A., Low, C.M.R., Perlin, K., Potapenko, A., Savy, P., Singh, S., Stecula, A., Thillaisundaram, A., Tong, C., Yakneen, S., Zhong, E.D., Zielinski, M., Žídek, A., Bapst, V., Kohli, P., Jaderberg, M., Hassabis, D., Jumper, J.M. Accurate Structure Prediction of Biomolecular Interactions with AlphaFold3. Nature 630, 493–500 (2024).

12. Krokidis, M.G., Koumadorakis, D.E., Lazaros, K., Ivantsik, O., Exarchos, T.P., Vrahatis, A.G., Kotsiantis, S., Vlamos, P. AlphaFold3: An Overview of Applications and Performance Insights. Int. J. Mol. Sci. 26, 3671 (2025).

13. Griffin, C., Howard, H., Paterson, A., Swanson, N., Bloxwich, D., Jumper, J., Kohli, P., Lundblad, N. Our approach to biosecurity for AlphaFold 3. Google DeepMind (2024).

14. Altschul, S.F., Madden, T.L., Schäffer, A.A., Zhang, J., Zhang, Z., Miller, W., Lipman, D.J. Gapped BLAST and PSI-BLAST: a new generation of protein database search programs. Nucleic Acids Res. 25, 3389–3402 (1997).

15. National Center for Biotechnology Information (NCBI). Protein BLAST. U.S. National Library of Medicine. Retrieved August 13, 2026, from https://blast.ncbi.nlm.nih.gov/Blast.cgi?PAGE=Protein (2026).

16. Ovchinnikov, S., Rettie, S., Favor, A., Abanades Kenyon, B., Batra, H., Amani, K. sokrypton/ColabDesign: v1.1.3. Zenodo. 10.5281/zenodo.15161108 (2025).

17. Pettersen, E.F., Goddard, T.D., Huang, C.C., Meng, E.C., Couch, G.S., Croll, T.I., Morris, J.H., Ferrin, T.E. UCSF ChimeraX: Structure Visualization for Researchers, Educators, and Developers. Protein Sci. 30, 70–82 (2021).

18. Wu, P., Chen, H., Coons, M., Crawford, T.D., Kirkpatrick, D.S., Murray, L., Ndubaku, C.O., Nonomiya, J., Pham, V., Schmidt, S., Smysczek, T., Vitorino, P., Ye, W., Harris, S.F. Structural Plasticity and Kinase Activation in a Cohort of MAP4K4 Structures. RCSB Protein Data Bank, PDB ID 4U40 (2016).

19. Abraham, M.J., Murtola, T., Schulz, R., Páll, S., Smith, J.C., Hess, B., Lindahl, E. GROMACS: High Performance Molecular Simulations through Multi-Level Parallelism from Laptops to Supercomputers. SoftwareX 1-2, 19–25 (2015).

20. Huang, J., Rauscher, S., Nawrocki, G., Ran, T., Feig, M., de Groot, B.L., Grubmüller, H., MacKerell, A.D. Jr. CHARMM36m: An Improved Force Field for Folded and Intrinsically Disordered Proteins. Nat. Methods 14, 71–73 (2017).

21. De Bondt, H.L., Rosenblatt, J., Jancarik, J., Jones, H.D., Morgan, D.O., Kim, S.-H. Crystal structure of cyclin-dependent kinase 2. Nature 363, 595–602 (1993).

22. Sharma, N., Naorem, L.D., Jain, S., Raghava, G.P.S. ToxinPred2: an improved method for predicting toxicity of proteins. Brief. Bioinform. 23, bbac174 (2022).

23. Sharma, N., Patiyal, S., Dhall, A., Pande, A., Arora, C., Raghava, G.P.S. AlgPred 2.0: an improved method for predicting allergenic proteins and mapping of IgE epitopes. Brief. Bioinform. 22, bbaa294 (2021).

24. Ozden, B., Cuesta Astroz, Y., Karaca, E. AlphaFold2 and AlphaFold3 lead to significantly different results in human-parasite interaction prediction. bioRxiv 2024.09.19.613643 (2024).

25. Strathern, M. “Improving ratings”: audit in the British university system. European Review 5, 305–321 (1997).

