## Supplementary Materials for "De Novo Design and AlphaFold3 Evaluation of Protein Binders Targeting Specific Sites of MAP4K4"

### Supplementary Material for De Novo Design and AlphaFold3 Evaluation of Protein Binders Targeting Specific Sites of MAP4K4

Table S1. Predicted biophysical and developability properties of the nine AlphaFold3-validated MAP4K4 binder candidates (Table 2), computed from each isolated binder chain's sequence (Section 3.10).

| Binder Name | AF3 ipTM | MW (Da) | pI | Instability index | ToxinPred2 score (call) | AlgPred2 ML / MERCI / Hybrid score (call) |
|---|---|---|---|---|---|---|
| BC-72 | 0.90 | 8464.7 | 5.18 | 65.73 | 0.42 (Non-Toxin) | 0.15 / 0.0 / 0.15 (Non-Allergen) |
| BC-128 | 0.89 | 14629.5 | 4.38 | 43.91 | 0.33 (Non-Toxin) | 0.22 / 0.0 / 0.22 (Non-Allergen) |
| BC-88 | 0.89 | 10740.6 | 10.67 | 47.38 | 0.44 (Non-Toxin) | 0.19 / 0.5 / 0.69 (Allergen) |
| PA2-june12 | 0.88 | 7955.2 | 5.54 | 63.54 | 0.67 (Toxin) | 0.19 / 0.0 / 0.19 (Non-Allergen) |
| BC-82 | 0.87 | 10255.3 | 6.13 | 74.23 | 0.42 (Non-Toxin) | 0.16 / 0.0 / 0.16 (Non-Allergen) |
| BC-53 | 0.84 | 6081.8 | 4.93 | 22.6 | 0.42 (Non-Toxin) | 0.29 / 0.0 / 0.29 (Non-Allergen) |
| BC-85 | 0.83 | 10648.9 | 9.39 | 44.22 | 0.40 (Non-Toxin) | 0.14 / 0.0 / 0.14 (Non-Allergen) |
| PA2-Des4num15 | 0.80 | 7970.0 | 4.81 | 52.43 | 0.76 (Toxin) | 0.27 / 0.0 / 0.27 (Non-Allergen) |
| BC-97 | 0.80 | 11378.0 | 8.29 | 44.04 | 0.57 (Non-Toxin) | 0.15 / 0.0 / 0.15 (Non-Allergen) |

Table S2. NCBI BLAST results for all 20 designed binder sequences (Section 3.7).

| **Binder Name** | **Closest BLASTp match by E-value** | **Accession number of closest match** | **E-value** | **Query Cover (%)** | **% Identity** |
|---|---|---|---|---|---|
| BC-72 | No matches returned | n/a | n/a | n/a | n/a |
| BC-128 | No matches returned | n/a | n/a | n/a | n/a |
| BC-88 | acyl-CoA dehydrogenase family protein *[Acidimicrobiales bacterium]* | MHB8670360.1 | 1.4 | 43 | 37 |
| PA2-june12 | No matches returned | n/a | n/a | n/a | n/a |
| BC-82 | uncharacterized protein *[Colletotrichum abscissum]* | XP_060404679.1 | 0.69 | 85 | 31 |
| BC-53 | SpoIID/LytB domain-containing protein *[Lachnotalea sp.]* | MHZ3560819.1 | 6.3 | 62 | 50 |
| BC-85 | DUF5060 domain-containing protein *[Anaerolineales bacterium]* | HEX6035026.1 | 8.4 | 98 | 28 |
| PA2-Des4num15 | No matches returned | n/a | n/a | n/a | n/a |
| BC-97 | hypothetical protein *[Micromonospora sp.* WMMA1363*]* | MDM4718081.1 | 1.7 | 63 | 33 |
| PA2-history | No matches returned | n/a | n/a | n/a | n/a |
| BC-50 | BamA/TamA family outer membrane protein *[Cyanobacteriota bacterium]* | MDY6938437.1 | 3.8 | 96 | 39 |
| PA2-des9n7 | No matches returned | n/a | n/a | n/a | n/a |
| BC-137 | EAL domain-containing protein *[Selenomonadales bacterium]* | MCI7478601.1 | 4.5 | 28 | 36 |
| BC-115 | No matches returned | n/a | n/a | n/a | n/a |
| BC-93 | beta-phosphoglucomutase family hydrolase *[Candidatus Gorgyraea atricola]* | MDP8229765.1 | 6.6 | 68 | 37 |
| BC-75 | No matches returned | n/a | n/a | n/a | n/a |
| BC-54 | carboxynorspermidine decarboxylase *[Alistipes sp.]* | WP_418426496.1 | 2.7 | 70 | 46 |
| BC-55 | No matches returned | n/a | n/a | n/a | n/a |
| BC-79 | No matches returned | n/a | n/a | n/a | n/a |
| BC-103 | tyrosine-type recombinase/integrase *[Staphylococcus pseudintermedius]* | MID7539303.1 | 7.8 | 67 | 30 |

Notes: E-values for all listed matches are well above the 0.05 significance threshold, consistent with coincidental similarities arising from querying sequences against a large database. BLAST settings are listed in Section 2.6.

### Supplementary Sequence 1: MAP4K4 target sequence submitted to AFS

Sequence of the MAP4K4 kinase-domain target chain submitted to the AlphaFold Server (AFS) in every job described in Section 2.5 of the main manuscript. This is a 295-residue subsequence of the full MAP4K4 kinase domain crystallization construct (PDB ID: 4U40), omitting the 17 N-terminal residues (GSANDSPAKSLVDIDLS) and the 20 C-terminal residues (KRGEKDETEYEYSGSEEGNS) that lie outside the ordered kinase-domain core.

```
>Supplementary_Sequence_1
SLRDPAGIFELVEVVGNGTYGQVYKGRHVKTGQLAAIKVMDVTEDEEEEIKLEINMLKKYSHHRNIATYY
GAFIKKSPPGHDDQLWLVMEFCGAGSITDLVKNTKGNTLKEDWIAYISREILRGLAHLHIHHVIHRDIKG
QNVLLTENAEVKLVDFGVSAQLDRTVGRRNTFIGTPYWMAPEVIACDENPDATYDYRSDLWSCGITAIEM
AEGAPPLCDMHPMRALFLIPRNPPPRLKSKKWSKKFFSFIEGCLVKNYMQRPSTEQLLKHPFIRDQPNER
QVRIQLKDHIDRTRK
```

---

### Supplementary Code 1: Auto-detect Binding Hotspots script

Python cell (run in the RFdiffusion/ColabDesign notebook, after the "Load RFdiffusion" cell) used to automatically score and rank candidate binding hotspots on the MAP4K4 target structure prior to binder design, as referenced in Section 2.2 of the main manuscript. The algorithm scores surface patches on four heuristics (surface geometry, hydrophobic/polar chemistry pattern, secondary-structure rigidity, and residue-cluster proximity) and outputs a ranked table plus a ready-to-use hotspot string.

```
#@title Auto-detect Binding Hotspots (run after "Load RFdiffusion")
# ─────────────────────────────────────────────────────────────────────────────
# Scores surface patches on your AlphaFold target using four heuristics:
#   H1 Geometry  - prefers grooves/knobs, rejects flat or ultra-narrow surfaces
#   H2 Chemistry - hydrophobic core surrounded by polar/charged rim
#   H3 Rigidity  - residues on helices/sheets score higher than loops
#   H4 Proximity - cluster of 3-5 residues all within 5 Å of each other
#
# Outputs a ranked table + a ready-to-paste `hotspot` string for the run cell.
# ─────────────────────────────────────────────────────────────────────────────

import subprocess, sys

# ── Install dependencies (silent) ─────────────────────────────────────────────
for pkg in ["freesasa", "biopython"]:
    subprocess.run([sys.executable, "-m", "pip", "install", "-q", pkg],
                   capture_output=True)

import os, re, warnings
import numpy as np
from itertools import combinations
warnings.filterwarnings("ignore")

from Bio import PDB
from Bio.PDB import DSSP
import freesasa

# ─────────────────────────────────────────────────────────────────────────────
#  USER SETTINGS  (edit these, then Run Cell)
# ─────────────────────────────────────────────────────────────────────────────
hotspot_pdb       = ""         #@param {type:"string"}
```

```
#@markdown 4-character PDB code (e.g. `4U40`) **or** leave blank to re-use the `pdb`
value already set in the run cell.
hotspot_chain     = "A"        #@param {type:"string"}
#@markdown Chain to search (usually A for single-chain AF structures).
top_n_results     = 5          #@param ["3","5","10"] {type:"raw"}
#@markdown How many candidate hotspots to display.
cluster_radius_A  = 5.0        #@param {type:"number"}
#@markdown Max Cα–Cα distance (Å) between any two residues in a cluster.
min_sasa_A2       = 10.0       #@param {type:"number"}
#@markdown Minimum solvent-accessible surface area (Ų) to consider a residue surface-
exposed.
auto_set_hotspot  = True       #@param {type:"boolean"}
#@markdown Automatically set the `hotspot` variable to the top-ranked cluster (ready
for the run cell).

# ──────────────────────────────────────────────────────────────────────
#  Residue chemistry dictionaries
# ──────────────────────────────────────────────────────────────────────
HYDROPHOBIC   = {"LEU","ILE","VAL","PHE","TRP","MET","TYR"}   # core "glue"
POLAR_CHARGED = {"ARG","LYS","ASP","GLU","ASN","GLN","HIS","SER","THR"}  # rim
"velcro"
HELIX_SS      = {"H","G","I"}   # α, 3₁₀, π helices
SHEET_SS      = {"E","B"}       # β-strand, β-bridge

# ──────────────────────────────────────────────────────────────────────
#  Helper: resolve the PDB file to analyze
# ──────────────────────────────────────────────────────────────────────
def _resolve_pdb(user_str):
    """Return a path to a .pdb file, using get_pdb() if needed."""
    if user_str and os.path.isfile(user_str):
        return user_str
    if "get_pdb" not in dir() and "get_pdb" not in globals():
        raise EnvironmentError(
            "Run the 'Load RFdiffusion' cell first so get_pdb() is available."
        )
    # Use the supplied code, or fall back to the `pdb` variable in the run cell
    code = user_str if user_str else globals().get("pdb", "")
    if not code:
        raise ValueError(
            "No PDB code provided. Either set `hotspot_pdb` below "
            "or run the RFdiffusion cell first so `pdb` is defined."
        )
    return get_pdb(code)

# ──────────────────────────────────────────────────────────────────────
#  Core scoring function
# ──────────────────────────────────────────────────────────────────────
def find_hotspots(pdb_file, chain_id="A",
                  top_n=5, cluster_dist=5.0, min_sasa=10.0):

    # ── Parse structure ─────────────────────────────────────────────
    parser = PDB.PDBParser(QUIET=True)
    structure = parser.get_structure("tgt", pdb_file)
    model = structure[0]
    chain = model[chain_id]
    residues = [r for r in chain if PDB.is_aa(r, standard=True) and "CA" in r]

    # ── DSSP: secondary structure ───────────────────────────────────
    dssp_map = {}
    try:
        dssp = DSSP(model, pdb_file, dssp="mkdssp")
        dssp_available = True
        for key in dssp.property_keys:
```

```
        pass  # just force load
    for res in residues:
        k = (chain_id, res.get_id())
        try:
            dssp_map[res.get_id()[1]] = dssp[k][2]   # SS code
        except KeyError:
            dssp_map[res.get_id()[1]] = "-"
except Exception:
    dssp_available = False
    for res in residues:
        dssp_map[res.get_id()[1]] = "?"

# ── freesasa: per-residue SASA ────────────────────────────────────────────
sasa_map = {}
try:
    fs_struct = freesasa.Structure(pdb_file)
    fs_result = freesasa.calc(fs_struct)
    for res in residues:
        rnum = res.get_id()[1]
        sel_name = f"res{rnum}"
        sel_str  = f"{sel_name}, (resi {rnum}) and (chain {chain_id})"
        area = freesasa.selectArea([sel_str], fs_struct, fs_result)
        sasa_map[rnum] = area.get(sel_name, 0.0)
except Exception as e:
    print(f"  ⚠ freesasa error ({e}); falling back to neighbour-count proxy.")
    # proxy: buried residues have many Cα neighbours
    all_ca = np.array([r["CA"].get_vector().get_array() for r in residues])
    for i, res in enumerate(residues):
        rnum = res.get_id()[1]
        dists = np.linalg.norm(all_ca - all_ca[i], axis=1)
        n_nb  = np.sum((dists > 0) & (dists < 8.0))
        sasa_map[rnum] = max(0.0, 80.0 - n_nb * 4.0)   # rough heuristic

# ── Build per-residue feature dict ───────────────────────────────────────
all_ca = np.array([r["CA"].get_vector().get_array() for r in residues])
res_nums = [r.get_id()[1] for r in residues]

data = {}
for i, res in enumerate(residues):
    rnum = res.get_id()[1]
    rname = res.get_resname()
    ca    = all_ca[i]
    sasa  = sasa_map.get(rnum, 0.0)
    ss    = dssp_map.get(rnum, "-")

    # H1 - Geometry: count Cα neighbours at 5 Å and 10 Å
    d_all  = np.linalg.norm(all_ca - ca, axis=1)
    n_5A   = int(np.sum((d_all > 0) & (d_all < 5.0)))
    n_10A  = int(np.sum((d_all > 0) & (d_all < 10.0)))
    # Groove/bowl fingerprint: moderate density at 10 Å, not packed at 5 Å
    # Flat < 6 at 10Å; buried > 18 at 10Å; slit: n_5A > 8
    if sasa < min_sasa or n_10A > 20 or n_5A > 8:
        geom = 0.0   # buried / too narrow
    elif n_10A < 5:
        geom = 0.2   # too flat
    else:
        ideal = 11
        geom  = max(0.0, 1.0 - abs(n_10A - ideal) / 9.0)

    # H2 - Chemistry
    if rname in HYDROPHOBIC and sasa >= min_sasa:
        chem = 1.0      # primary glue candidate
    elif rname in POLAR_CHARGED and sasa >= min_sasa:
```

```
        chem = 0.55     # rim candidate
    else:
        chem = 0.1

    # H3 - Rigidity
    if ss in HELIX_SS:
        rigid = 1.0
    elif ss in SHEET_SS:
        rigid = 0.90
    elif ss == "?":
        rigid = 0.5     # unknown → neutral
    else:
        rigid = 0.05    # loop / coil

    # pLDDT bonus (AF stores pLDDT in B-factor)
    try:
        plddt = res["CA"].get_bfactor()
    except Exception:
        plddt = 70.0
    plddt_score = min(1.0, max(0.0, (plddt - 50.0) / 50.0))

    single = (geom * 0.35 + chem * 0.35 + rigid * 0.20 + plddt_score * 0.10)

    data[rnum] = dict(
        resname=rname, coord=ca, sasa=sasa, ss=ss,
        geom=geom, chem=chem, rigid=rigid, plddt=plddt,
        single=single,
    )

# ── H4 - Cluster enumeration and scoring ─────────────────────────────────
# Only consider surface-exposed residues with non-trivial single score
candidates_res = [r for r in res_nums
                  if data[r]["sasa"] >= min_sasa and data[r]["single"] > 0.08]

coord_arr = {r: data[r]["coord"] for r in candidates_res}

clusters = []
seen_frozensets = set()

# Greedy seeded expansion: for each residue, grow cluster greedily
for seed in candidates_res:
    c_vec = coord_arr[seed]
    # All neighbours within cluster_dist
    nbrs = [(np.linalg.norm(coord_arr[r] - c_vec), r)
            for r in candidates_res if r != seed
            and np.linalg.norm(coord_arr[r] - c_vec) <= cluster_dist]
    nbrs.sort()

    for size in range(2, 5):   # cluster = seed + 2..4 neighbours → 3..5 total
        if len(nbrs) < size:
            break
        members = [seed] + [r for _, r in nbrs[:size]]
        fs = frozenset(members)
        if fs in seen_frozensets:
            continue
        seen_frozensets.add(fs)

        # Verify max pairwise distance (tight cluster requirement)
        coords_m = np.array([coord_arr[r] for r in members])
        max_pd = max(
            np.linalg.norm(coords_m[ii] - coords_m[jj])
            for ii, jj in combinations(range(len(members)), 2)
        )
```

```
            if max_pd > cluster_dist * 1.6:
                continue

            rnames = [data[r]["resname"] for r in members]
            n_hyd  = sum(1 for rn in rnames if rn in HYDROPHOBIC)
            n_pol  = sum(1 for rn in rnames if rn in POLAR_CHARGED)

            # Must have at least 1 hydrophobic core residue
            if n_hyd == 0:
                continue

            # Chemistry pattern score: ~30-50% hydrophobic is ideal
            ratio = n_hyd / len(members)
            chem_pat = max(0.0, 1.0 - abs(ratio - 0.40) * 2.5)
            if n_pol == 0:
                chem_pat *= 0.5   # penalise missing polar rim

            avg_geom  = np.mean([data[r]["geom"]   for r in members])
            avg_rigid = np.mean([data[r]["rigid"]  for r in members])
            avg_plddt = np.mean([data[r]["plddt"]  for r in members])

            # Size bonus: 4-5 residues preferred
            size_bonus = 0.05 if len(members) >= 4 else 0.0

            proximity = max(0.0, 1.0 - max_pd / (cluster_dist * 1.6))

            score = (avg_geom  * 0.30 +
                     chem_pat  * 0.30 +
                     avg_rigid * 0.20 +
                     proximity * 0.15 +
                     min(1.0, avg_plddt / 100.0) * 0.05 +
                     size_bonus)

            hotspot_str = ", ".join(f"{chain_id}{r}" for r in sorted(members))

            clusters.append(dict(
                members=sorted(members),
                hotspot_str=hotspot_str,
                score=score,
                n_hyd=n_hyd, n_pol=n_pol,
                max_dist=max_pd,
                avg_rigid=avg_rigid,
                avg_geom=avg_geom,
                chem_pat=chem_pat,
                avg_plddt=avg_plddt,
                rnames=rnames,
            ))

    if not clusters:
        print("⚠  No clusters found. Try increasing cluster_radius_A or lowering
min_sasa_A2.")
        return [], data

    # Deduplicate: keep clusters whose residues overlap <50% with any higher-scored
cluster
    clusters.sort(key=lambda x: -x["score"])
    final = []
    for c in clusters:
        cs = frozenset(c["members"])
        if all(len(cs & frozenset(f["members"])) / len(cs) <= 0.5 for f in final):
            final.append(c)
        if len(final) >= top_n:
```

```
            break

    return final, data

# ─────────────────────────────────────────────────────────────────────────────
#  Run analysis
# ─────────────────────────────────────────────────────────────────────────────
print("=" * 70)
print("  Binding Hotspot Auto-Detection")
print("=" * 70)

target_pdb = _resolve_pdb(hotspot_pdb)
print(f"\n  Target structure : {target_pdb}")
print(f"  Chain            : {hotspot_chain}")
print(f"  Cluster radius   : {cluster_radius_A} Å")
print(f"  Min SASA         : {min_sasa_A2} Ų")
print("\n  Scoring residues ... (this takes ~10-30 seconds)\n")

results, res_data = find_hotspots(
    target_pdb,
    chain_id    = hotspot_chain,
    top_n       = top_n_results,
    cluster_dist= cluster_radius_A,
    min_sasa    = min_sasa_A2,
)

if not results:
    print("No candidates found. See warning above.")
else:
    # ── Pretty table ─────────────────────────────────────────────────────────
    HDR = f"  {'Rank':<5} {'Score':>6}  {'Residues (hotspot string)':<34}  {'Hyd':>4}
{'Pol':>4}  {'Geom':>5} {'Rigid':>6} {'Chem':>5}  {'pLDDT':>6}  {'MaxDist':>8}"
    print(HDR)
    print("  " + "─" * (len(HDR) - 2))

    for rank, c in enumerate(results, 1):
        label = "← TOP PICK ✓" if rank == 1 else ""
        print(f"  {rank:<5} {c['score']:>6.3f}  {c['hotspot_str']:<34}  "
              f"{c['n_hyd']:>4} {c['n_pol']:>4}  "
              f"{c['avg_geom']:>5.2f} {c['avg_rigid']:>6.2f} {c['chem_pat']:>5.2f}  "
              f"{c['avg_plddt']:>6.1f}  {c['max_dist']:>7.1f}Å  {label}")

    print()
    print("  Column guide:")
    print("    Score   - composite hotspot score (higher = better)")
    print("    Hyd/Pol - hydrophobic / polar+charged residue count in cluster")
    print("    Geom    - H1 geometry (groove/knob) score [0-1]")
    print("    Rigid   - H3 secondary-structure rigidity score [0-1]")
    print("    Chem    - H2 chemistry pattern score [0-1]")
    print("    pLDDT   - mean AlphaFold confidence (>70 recommended)")
    print("    MaxDist - largest pairwise Cα distance in the cluster")
    print()

    top = results[0]
    print("─" * 70)
    print(f"  TOP CANDIDATE  →  hotspot = \"{top['hotspot_str']}\"")
    print("─" * 70)

    # Residue breakdown for top candidate
    print(f"\n  Residue breakdown (top cluster):")
    for r in top["members"]:
        d   = res_data[r]
        tag = ("HYDROPHOBIC" if d["resname"] in HYDROPHOBIC else
```

```
                "POLAR/CHG"  if d["resname"] in POLAR_CHARGED else "OTHER")
        ss_label = ("helix" if d["ss"] in {"H","G","I"} else
                    "sheet" if d["ss"] in {"E","B"} else
                    "loop " if d["ss"] in {"-","T","S"} else "?    ")
        print(f"    {hotspot_chain}{r:>4}  {d['resname']:<4}  {tag:<12}  "
              f"{ss_label}  SASA={d['sasa']:>6.1f} Ų  pLDDT={d['plddt']:>5.1f}")

    # ── Optionally set the global `hotspot` variable ──────────────────────────
    if auto_set_hotspot:
        hotspot = top["hotspot_str"]
        print(f"\n  ✅  `hotspot` variable has been set to: \"{hotspot}\"")
        print("     Run the RFdiffusion cell directly — no copy-paste needed.")
    else:
        print(f"\n  Copy this into the `hotspot` field of the run cell:")
        print(f"    {top['hotspot_str']}")

print("\n" + "=" * 70)
```
